# PERK Signaling Promotes Mitochondrial Elongation By Remodeling Membrane Phosphatidic Acid

**DOI:** 10.1101/2022.02.23.481593

**Authors:** Valerie Perea, Christian Cole, Justine Lebeau, Vivian Dolina, Kelsey R. Baron, Aparajita Madhavan, Jeffery W. Kelly, Danielle A. Grotjahn, R. Luke Wiseman

**Author notes:** These authors contributed equally (Christian Cole, Justine Lebeau). To whom correspondences should be addressed: R. Luke Wiseman, Department of Molecular Medicine, Scripps Research, La Jolla, CA 92037.

## Abstract

Endoplasmic reticulum (ER) stress and mitochondrial dysfunction are linked in the onset and pathogenesis of numerous diseases. This has led to considerable interest in defining the mechanisms responsible for regulating mitochondria during ER stress. The PERK signaling arm of the unfolded protein response (UPR) has emerged as a prominent ER stress-responsive signaling pathway that regulates diverse aspects of mitochondrial biology. Here, we show that PERK activity also promotes adaptive remodeling of mitochondrial membrane phosphatidic acid (PA) to induce protective mitochondrial elongation during ER stress. We find that PERK activity is required for ER stress-dependent increases in both cellular PA and YME1L-dependent degradation of the intramitochondrial PA transporter PRELID1. Together, these processes lead to the accumulation of PA on the outer mitochondrial membrane where it induces mitochondrial elongation by inhibiting mitochondrial fission. Our results establish a new role for PERK in the adaptive remodeling of mitochondrial phospholipids and demonstrate that PERK-dependent PA regulation functions to adapt organellar shape in response to ER stress.

## INTRODUCTION

Endoplasmic reticulum (ER) and mitochondrial function are coordinated through the inter-organellar transport of metabolites such as lipids and Ca^2+^.^1–3^ As a consequence of this coordination, ER stress can be transmitted to mitochondria and promote mitochondrial dysfunctions implicated in the pathophysiology of numerous diseases including diabetes, cardiovascular disorders, and many neurodegenerative diseases.^4–16^ This pathologic relationship between ER stress and mitochondria has led to significant interest in identifying the stress-responsive signaling pathways responsible for regulating mitochondria in response to ER insults.

The PERK arm of the unfolded protein response (UPR) has emerged as a prominent stress-responsive signaling pathway involved in regulating mitochondria during ER stress.^17–19^ PERK is an ER transmembrane protein that is activated in response to ER stress through a mechanism involving oligomerization and autophosphorylation of its cytosolic kinase domain (**Fig. 1A**).^20–22^ Activated PERK selectively phosphorylates serine 51 of the α subunit of eukaryotic initiation factor 2 (eIF2α). Phosphorylated eIF2α prevents formation of ribosomal initiation complexes leading to global mRNA translational attenuation, which functions to reduce the load of newly synthesized proteins during ER stress.^20–22^ PERK-dependent eIF2α phosphorylation also leads to the selective translation and activation of transcription factors, such as ATF4, through upstream open reading frames (uORFs) in the promoters of these genes.^20–23^ ATF4 regulates the expression of several stress-responsive genes including redox factors, amino acid biosynthesis genes, the eIF2α phosphatase *PPP1R15A*/*GADD34*, and the pro-apoptotic transcription factor *DDIT3*/*CHOP*.^23–25^ Through this combination of translational attenuation and transcriptional signaling, PERK promotes both adaptive and pro-apoptotic signaling in response to varying levels and extents of ER stress.^20–31^

**Figure 1.**
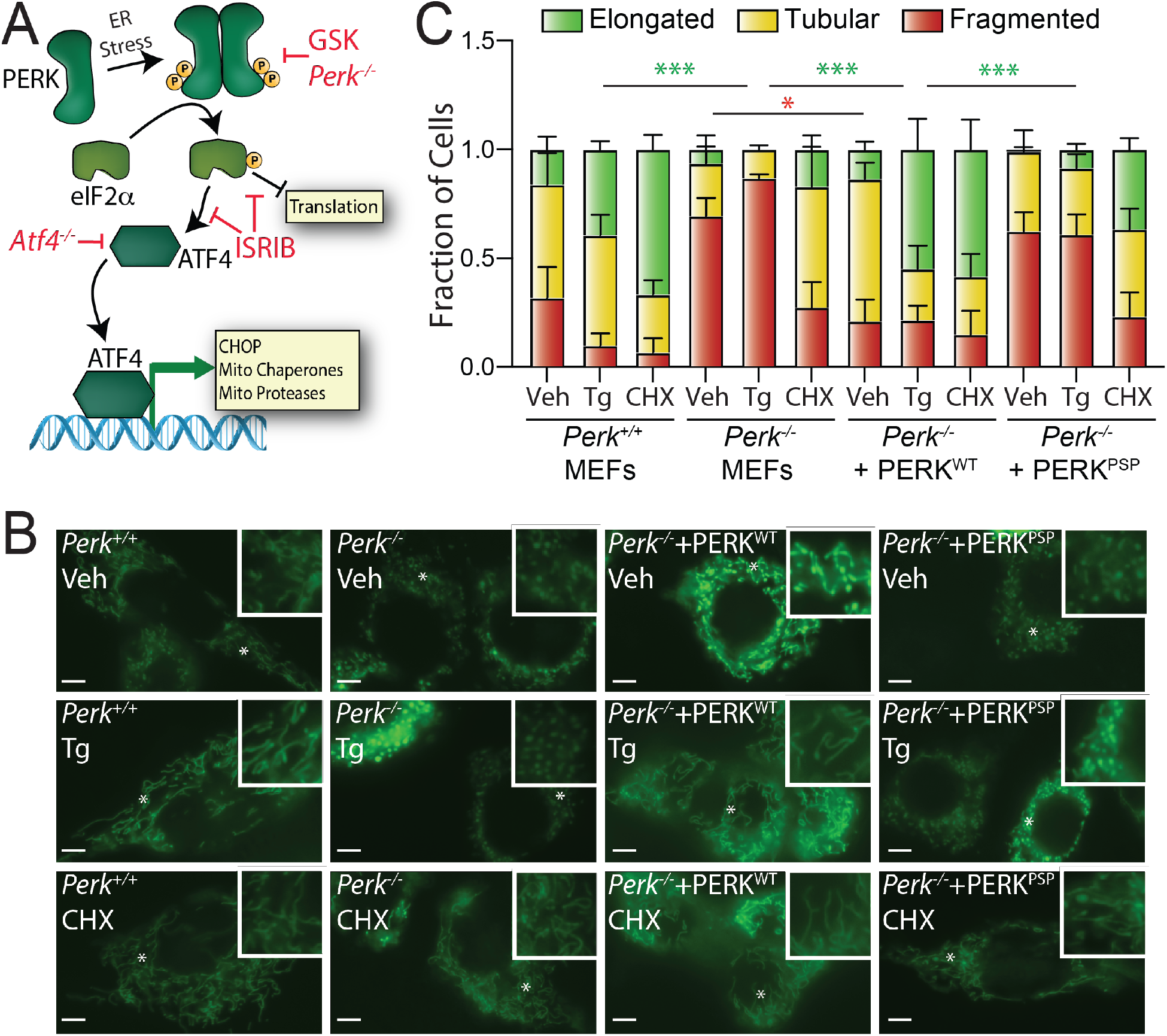
ER stress induced mitochondrial elongation is impaired in cells expressing hypomorphic PERK variants. **A.** Illustration showing the mechanism of PERK-regulated transcriptional and translational signaling. Specific genetic and pharmacologic manipulations used to disrupt PERK signaling are shown. **B.** Representative images of *Perk^+/+^* MEFs, *Perk^−/−^* MEFs, or *Perk^−/−^* MEFs transfected with wild-type PERK^WT^ or the PSP-associated PERK allele (PERK^PSP^) treated for 6 h with thapsigargin (Tg; 500 nM) or cycloheximide (CHX; 50 μg/mL). The inset shows 2-fold magnification of the image centered on the asterisk. Scale bars, 5 μm. **C**. Quantification of fragmented (red), tubular (yellow), or elongated (green) mitochondria from the images shown in (**B**). Error bars show SEM for n=3 independent experiments. *p<0.05, ***p<0.005 for 2-way ANOVA (red indicates comparison between fragmented mitochondria fractions; green indicates comparisons between elongated mitochondria fractions).

PERK localizes to ER-mitochondrial contact sites, positioning this protein to coordinate regulation of these two organelles in response to cellular insults.^32^ Consistent with this, PERK signaling regulates diverse aspects of mitochondrial proteostasis and function in response to stress. PERK regulates mitochondrial protein import, biogenesis, and cristae remodeling in brown adipocytes in response to cold exposure or beta-adrenergic stimulation.^33,34^ Further, the PERK-regulated transcription factor ATF4 increases mitochondrial respiratory chain activity during ER stress or nutrient deprivation through a mechanism involving SCAF1-dependent increases in supercomplex formation.^35^ ATF4 also regulates the expression of numerous mitochondrial proteostasis factors including the mitochondrial HSP70 *HSPA9* and the AAA+ quality control protease *LONP1* to increase mitochondrial proteostasis capacity during ER stress.^25,36^ Further, PERK-dependent translational attenuation regulates mitochondrial protein import by selectively decreasing protein concentrations of the core TIM23 subunit Tim17A, a process dependent on the mitochondrial AAA+ protease YME1L.^37^

PERK signaling also promotes mitochondrial elongation downstream of eIF2α phosphorylation-dependent translational attenuation.^38^ This increase in mitochondrial length functions to protect mitochondria during ER stress by preventing premature fragmentation and regulating mitochondrial respiratory chain activity.^38^ However, the mechanistic basis of PERK-dependent mitochondrial elongation was previously undefined. Here, we show that PERK induces mitochondrial elongation through the adaptive remodeling of mitochondrial membrane phosphatidic acid (PA). Our results suggest a model whereby PERK signaling both increases total mitochondrial PA and inhibits trafficking of PA to the inner mitochondrial membrane. This leads to the accumulation of PA on the outer mitochondrial membrane where it induces mitochondrial elongation by inhibiting mitochondrial fission. These results define a new role for PERK in regulating the amount and localization of mitochondrial membrane phospholipids and show that this remodeling is important for adapting mitochondrial morphology during ER stress.

## RESULTS

### Hypomorphic PERK variants inhibit ER stress induced mitochondrial elongation

Pharmacologic inhibition of PERK signaling blocks mitochondrial elongation induced by ER stress.^38^ Here, we further probed the dependence of ER stress induced mitochondrial elongation on PERK activity in *Perk^−/−^* MEFs.^24^ We transfected *Perk^+/+^* or *Perk^−/−^* MEFs with mitochondrial targeted GFP (^mt^GFP) and monitored mitochondrial morphology in cells treated with either the ER stressor thapsigargin (Tg; a SERCA inhibitor) or the translation inhibitor cycloheximide (CHX) – the latter of which induces mitochondrial elongation independent of PERK signaling.^38,39^ *Perk^−/−^* MEFs showed increases in fragmented mitochondria in the absence of treatment (**Fig. 1B,C**). Tg-induced mitochondrial elongation was also impaired in *Perk*-deficient cells. However, CHX treatment reduced mitochondrial fragmentation in *Perk*-deficient cells, indicating that these cells are not deficient in their ability to increase length in response to reduced translation. Reconstitution of *Perk^−/−^* MEFs with wild-type PERK restored basal mitochondrial morphology and rescued Tg-induced mitochondrial elongation. However, a hypomorphic PERK haplotype implicated in progressive supranuclear palsy (PSP; PERK^PSP^)^40,41^ did not significantly impact basally fragmented mitochondria or rescue Tg-induced mitochondrial elongation in *Perk*-deficient cells. We confirmed similar expression of PERK^WT^ and PERK^PSP^ in *Perk^−/−^* MEFs by immunoblotting (**Fig. S1**). These results further implicate PERK signaling in ER stress induced mitochondrial elongation and demonstrate that genetic disruptions in PERK signaling impair the regulation of mitochondrial morphology in the presence or absence of ER stress.

### Overexpression of cytosolic PA lipases inhibits ER stress induced mitochondrial elongation

Mitochondrial morphology is defined by the relative activities of GTPases localized to the inner and outer mitochondrial membranes that regulate organellar fission and fusion. These include the pro-fission GTPase DRP1 of the outer mitochondrial membrane (OMM) and the pro-fusion GTPases MFN1 and MFN2 of the OMM and OPA1 of the inner mitochondrial membrane (IMM).^42–46^ Stress-induced changes in mitochondrial shape are primarily dictated through posttranslational regulation of these GTPases to alter the relative activities of fusion and fission pathways.^42–46^ Previous results indicate that PERK signaling does not significantly influence the posttranslational regulation of these GTPases^38^, suggesting that ER stress induced mitochondrial elongation proceeds through an alternative mechanism.

Mitochondrial elongation can be induced by the accumulation of saturated PA on the OMM through mechanisms including PA-dependent inhibition of the pro-fission GTPase DRP1.^47–51^ Interestingly, PERK was previously shown to increase cellular PA during ER stress through a mechanism dependent on PERK kinase activity, but not signaling downstream of eIF2α phosphorylation.^52^ We confirmed that treatment with Tg increases PA in mitochondria-enriched fractions and whole cell extracts from MEF cells measured by mass spectrometry, biochemical assays, and ELISA (**Fig. 2A,B**, **Fig. S2A,D-E**). Another phospholipid, phosphatidylcholine (PC) was not affected in enriched mitochondria (**Fig. 2A**). Similar results were observed in HeLa cells (**Fig. S2B,C**). Co-treatment with the PERK inhibitor GSK2656157, a compound that directly inhibits PERK kinase activity (**Fig. 1A**)^53^, reduced Tg-dependent increases of PA in both MEF and HeLa cells (**Fig. 2B**, **Fig. S2A-C**). This indicates that ER stress-dependent increases in PA requires PERK kinase activity, as previously reported.^52^ However, co-treatment of MEF cells with Tg and ISRIB, a compound that blocks PERK signaling downstream of eIF2α phosphorylation (**Fig. 1A**)^54^, did not appear to mitigate ER stress induced PA increases in mitochondria enriched fractions or whole cell extracts (**Fig. S2D,E**). This is consistent with previous results suggesting that ER stress increases PA through a mechanism selectively dependent on PERK kinase activity, relative to signaling downstream of eIF2α phosphorylation.^52^

**Figure 2.**
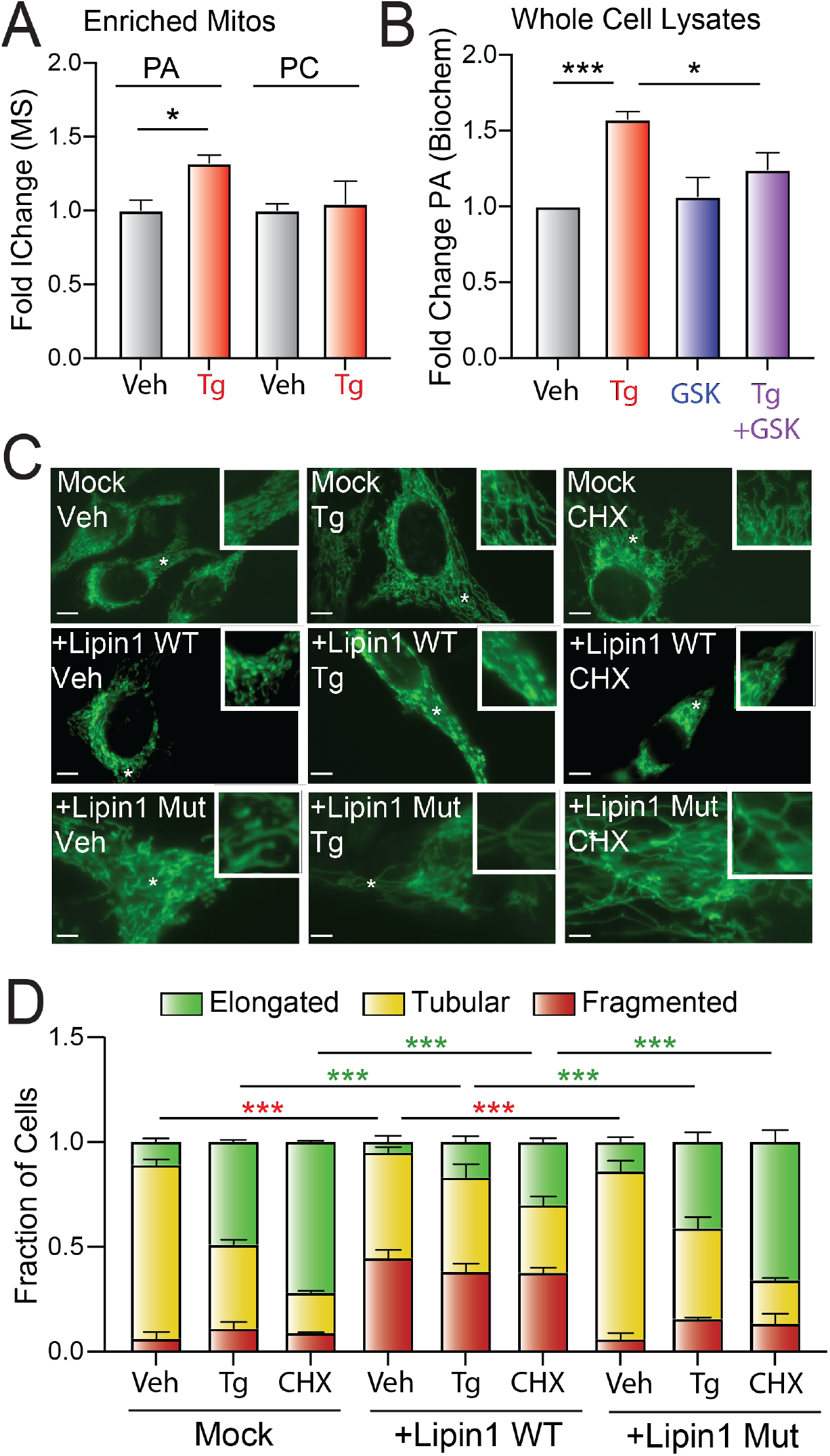
Overexpression of PA lipases inhibits ER stress induced mitochondrial elongation. **A.** Relative amounts of phosphatidic acid (PA) and phosphatidylcholine (PC), measured by mass spectrometry, in mitochondrial fractions isolated from MEF cells treated for 6 h with vehicle or thapsigargin (Tg; 500 nM). Error bars show SEM for n=3 replicates. *p<0.05 for unpaired t-test. **B**. Relative amounts of phosphatidic acid (PA), measured by biochemical assay (see Materials and Methods), in whole cell lysates prepared from MEF cells treated for 3 h with vehicle or thapsigargin (Tg; 500 nM). Error bars show SEM for n=3 replicates. *p<0.05, ***p<0.005 for paired t-test. **C.** Representative images of HeLa cells expressing ^mt^GFP transfected with mock, wild-type Lipin1 (Lipin1^wt^), or catalytically inactive Lipin1 (Lipin1^mut^) treated for 3 h with thapsigargin (Tg; 500 nM) or cycloheximide (CHX; 50 μg/mL). The inset shows 2-fold magnification of the image centered on the asterisk. Scale bars, 5 μm. **D.** Quantification of fragmented (red), tubular (yellow), or elongated (green) mitochondria from the images shown in (**D**). Error bars show SEM for n=3 experiments. p-value reflects comparisons of elongated (green) or fragmented (red) mitochondria populations for the indicated conditions. ***p<0.005 for 2-way ANOVA (red indicates comparison between fragmented mitochondria fractions; green indicates comparisons between elongated mitochondria fractions).

We next determined the dependence of PERK-regulated mitochondrial elongation on PA by monitoring mitochondrial morphology in Tg-treated HeLa cells co-overexpressing ^mt^GFP and Lipin1 – a cytosolic PA lipase that converts PA to diacylglycerol (DAG).^48,49,55,56^ We confirmed that Lipin1 overexpression reduced cellular PA (**Fig. S2F**). Overexpression of wild-type, but not catalytically inactive, Lipin1 increased basal mitochondrial fragmentation and inhibited Tg-induced mitochondrial elongation (**Fig 2C,D**). Similar results were observed in cells treated with CHX. Importantly, Lipin1 overexpression did not significantly influence increases of ATF4 protein, the expression of ATF4 target genes (e.g., *Asns*, *Chop*), or TIM17A degradation in Tg-treated cells (**Fig. S2G,H**). This indicates that Lipin1 overexpression did not influence upstream PERK signaling (**Fig. 1A**), but instead inhibited ER stress induced mitochondrial elongation at a step downstream of PERK-dependent eIF2α phosphorylation. Overexpression of PA-PLA1 – a cytosolic lipase that converts PA to lysophosphatidic acid^49^ – similarly inhibited mitochondrial elongation in cells treated with Tg or CHX without impacting other aspects of PERK signaling (**Fig. S2I-L**). Collectively, these results indicate that genetic depletion of PA blocks ER stress induced mitochondrial elongation, implicating PA in this process.

### ER stress prevents DRP1-dependent mitochondrial fragmentation

Accumulation of PA on the OMM promotes mitochondrial elongation by inhibiting the pro-fission GTPase DRP1.^51^ This was previously demonstrated by showing that genetically increasing PA on the OMM by overexpressing mitoPLD – an OMM lipase that converts cardiolipin to PA – basally increased mitochondrial elongation and inhibited DRP1-dependent mitochondrial fragmentation induced by the uncoupler carbonyl cyanide m-chlorophenylhydrazone (CCCP).^51,57^ Consistent with this, we confirmed that mitoPLD overexpression in HeLa cells increased basal mitochondrial elongation and inhibited CCCP-induced mitochondrial fragmentation (**Fig. S3A,B**). Pretreatment with Tg similarly reduced CCCP-induced mitochondrial fragmentation. However, Tg-pretreatment also blocked CCCP-induced proteolytic cleavage of the inner membrane GTPase OPA1 (**Fig. S3C**) – a biological process upstream of DRP1-dependent mitochondrial fragmentation induced by membrane uncoupling.^42–46,58^ This appears to result from Tg-dependent increases in mitochondrial membrane polarity, which prevent efficient mitochondrial uncoupling in CCCP-treated cells (**Fig. S3D**). Regardless, the Tg-dependent inhibition of OPA1 processing in cells treated with CCCP precluded our ability to determine whether ER stress inhibits DRP1 activity under these conditions.

To circumvent this problem, we monitored mitochondria morphology in MEF^mtGFP^ cells pretreated with Tg and then challenged with ionomycin – a Ca^2+^ ionophore that increases cytosolic Ca^2+^.^42–46,59^ Increases in cytosolic Ca^2+^ induced by short (<30 min) treatments with ionomycin or Tg promotes DRP1-dependent mitochondrial fragmentation through a mechanism independent of membrane uncoupling or OPA1 processing.^59^ However, pretreatment for 3 h with Tg – a timepoint sufficient to increase PA and induce mitochondrial elongation – inhibits ionomycin-induced mitochondrial fragmentation (**Fig. 3A,B**). This inhibition is reversed by co-treatment with ISRIB, a small molecule that blocks eIF2α phosphorylation-dependent signaling downstream of PERK (**Fig. 1A**), confirming that this effect can be attributed to PERK signaling and not dysregulation of intracellular Ca^2+^ induced by the combined treatment of Tg and ionomycin. These results are consistent with a model whereby ER stress promotes mitochondrial elongation through a mechanism involving inhibition of DRP1 activity, as was reported previously for mitoPLD overexpression ^51^.

**Figure 3.**
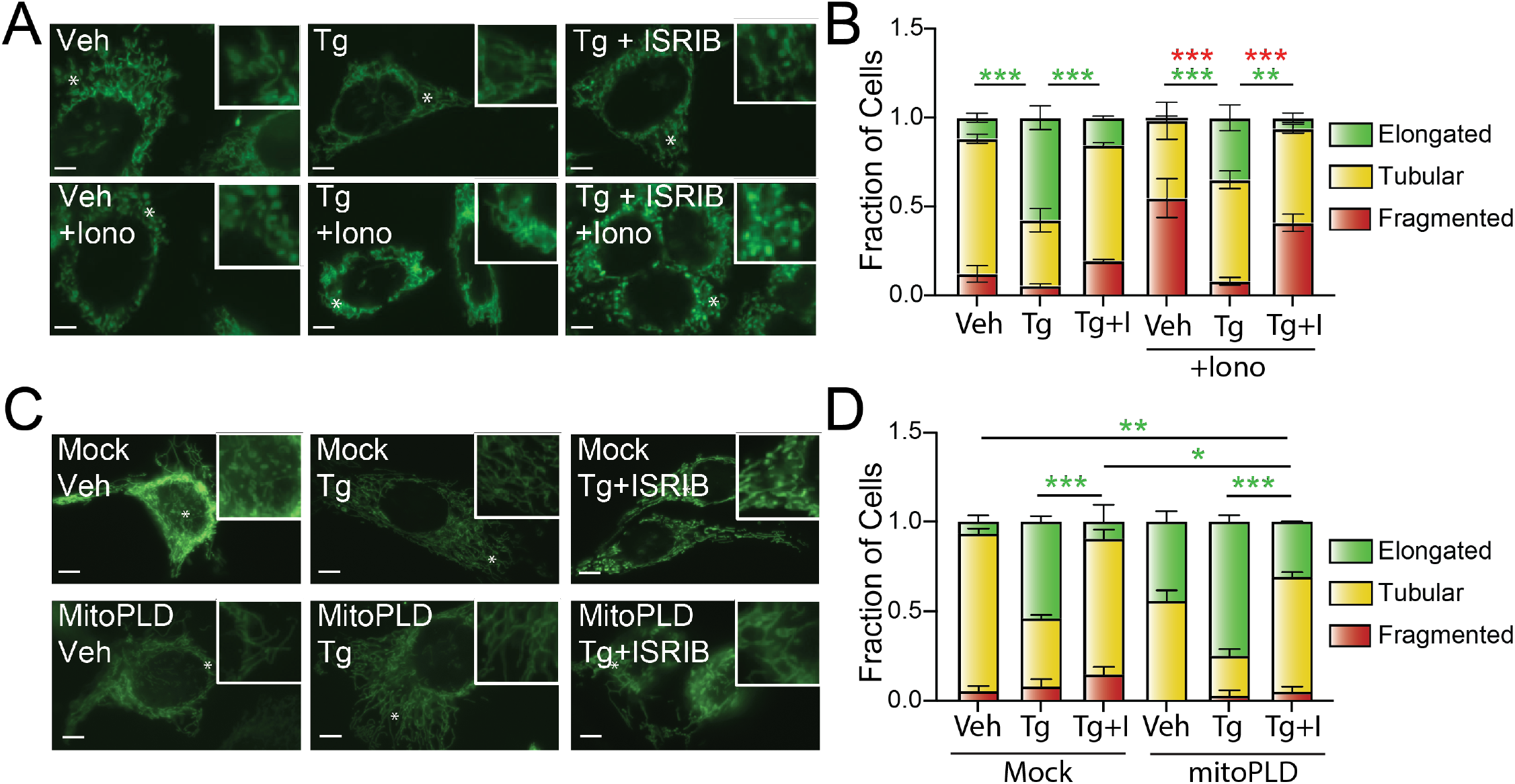
ER stress induced mitochondrial elongation inhibits Ionomycin-induced mitochondrial fragmentation. **A.** Representative images of MEF^mtGFP^ cells pre-treated for 3 h with vehicle, thapsigargin (Tg; 500 nM), or Tg and ISRIB (0.2 μM) then challenged with vehicle or ionomycin (Iono; 1 μM) for 30 min. The inset shows 2-fold magnification of the image centered on the asterisk. Scale bars, 5 μm. **B.** Quantification of fragmented (red), tubular (yellow), or elongated (green) mitochondria from the images shown in (**A**). Error bars show SEM for n=3 experiments. **p<0.01, ***p<0.005 for 2-way ANOVA (red indicates comparison between fragmented mitochondria fractions; green indicates comparisons between elongated mitochondria fractions). **C.** Representative images of HeLa cells expressing ^mt^GFP transfected with mock or mitoPLD^GFP^ and then treated with vehicle, Tg (500 nM), or Tg + ISRIB (0.2 μM). The inset shows 2-fold magnification of the image centered on the asterisk. Scale bars, 5 μm. Note the expression of mitoPLD^GFP^ did not impair our ability to accurately monitor mitochondrial morphology in these cells. **D.** Quantification of fragmented (red), tubular (yellow), or elongated (green) mitochondria from the images shown in (**C**). Error bars show SEM for n=3 experiments. p-value reflects comparisons of elongated (green) mitochondria populations for the indicated conditions. *p<0.05, **p<0.01, ***p<0.005 for 2-way ANOVA (green indicates comparisons between elongated mitochondria fractions).

### Mitochondrial elongation induced by mitoPLD overexpression is further enhanced by ER stress

Overexpression of mitoPLD increases cellular PA to similar extents to that observed in Tg-treated cells (**Fig. S3E**). This provides an opportunity to determine whether increases in PA are sufficient to induce the mitochondrial elongation observed during ER stress. To test this, we monitored mitochondrial morphology in HeLa cells overexpressing mitoPLD and treated with or without Tg. Despite similar increases in PA (**Fig. S3E**), mitoPLD overexpression did not increase mitochondrial elongation to the same extent observed upon Tg treatment (**Fig. 3C,D**). Further, treatment with Tg increased mitochondrial elongation in mitoPLD overexpressing cells, as compared to untreated controls (**Fig. 3C,D**). This increase was blocked by co-treatment with the PERK signaling inhibitor ISRIB, indicating that this enhanced elongation could be attributed to PERK signaling downstream of eIF2α phosphorylation (**Figs. 1A**, **3C,D**). Overexpression of mitoPLD did not significantly impact other aspects of PERK signaling in Tg-treated cells such as TIM17A degradation (**Fig. S3F**), suggesting that this effect cannot be attributed to altered or increased PERK activity. These results suggest that PERK-dependent increases in PA, on its own, may not be sufficient to explain the increase in mitochondrial elongation observed during ER stress and that other PERK-regulated activities induced downstream of eIF2α phosphorylation are also likely involved in this process.

### PERK signaling leads to reductions in the intramitochondrial PA transporter PRELID1 during ER stress

ER stress induced mitochondrial elongation is inhibited by shRNA-depletion of *YME1L* in HeLa cells.^38^ This suggests a potential role for YME1L in this process. We further confirmed the dependence of Tg-induced mitochondrial elongation on YME1L in MEF^mtGFP^ cells where *Yme1l* was deleted by CRISPR (**Fig. S4A,B**). PRELID1, an intermembrane space protein that transports PA from the OMM to the IMM (**Fig. 4A**)^55,56^, is a known substrate of YME1L.^60–62^ Interestingly, PRELID1 is a short-lived protein whose levels are highly sensitive to translational attenuation.^63^ Consistent with this, PRELID1 levels are rapidly reduced in MEF^mtGFP^ cells treated with CHX (**Fig. S4C**). This CHX-dependent reduction in PRELID1 was blocked in *Yme1l*-deficient cells (**Fig. 4B**), confirming that PRELID1 is degraded by YME1L under these conditions. Identical results were observed for TIM17A, another short-lived mitochondrial protein degraded by YME1L downstream of translation inhibition (**Fig. 4B** and **Fig. S4C**).^37^

**Figure 4.**
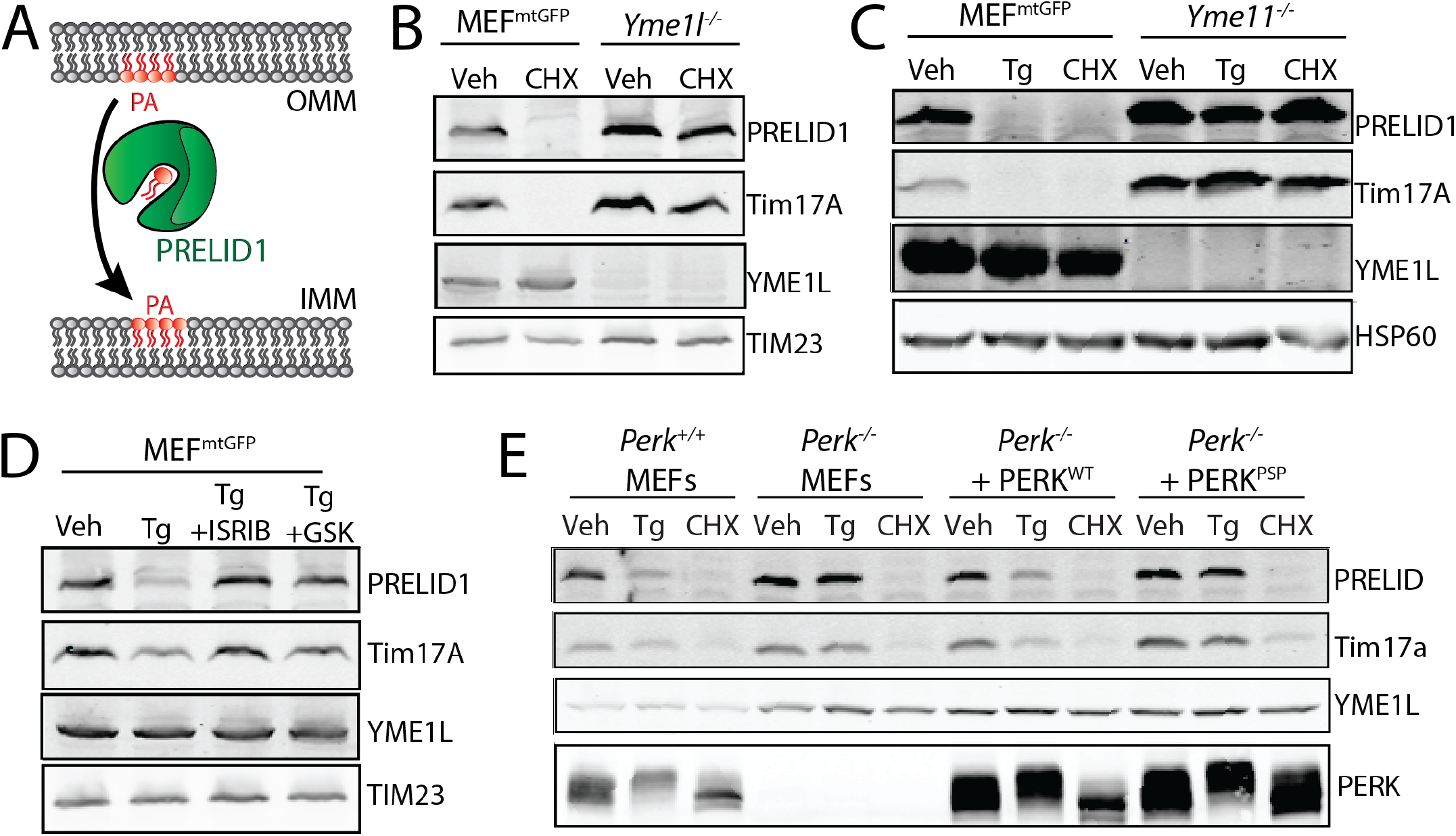
ER stress reduces PRELID1 through a YME1L-dependent mechanism downstream of PERK-dependent translational attenuation. **A.** Illustration showing the PRELID1-dependent trafficking of PA from the outer to inner mitochondrial membranes (OMM and IMM, respectively). **B**. Immunoblot of lysates prepared from MEF^mtGFP^ cells and *Yme1l*-deficient MEF^mtGFP^ cells treated for 3 h with vehicle or cycloheximide (CHX; 50 μg/mL). **C**. Immunoblot of lysates prepared from MEF^mtGFP^ cells and *Yme1l*-deficient MEF^mtGFP^ cells treated for 3 h with vehicle, thapsigargin (Tg; 500 nM) or cycloheximide (CHX; 50 μg/mL). **D.** Immunoblot of MEF^mtGFP^ cells treated with vehicle, Tg (500 nM), Tg and ISRIB (0.2 μM), or Tg and GSK2656157 (GSK; 1 μM). **E**. Immunoblot of lysates prepared from *Perk^+/+^* MEFs, *Perk^−/−^* MEFs, or *Perk^−/−^* MEFs transfected with wild-type PERK^WT^ or the PSP-associated PERK allele (PERK^PSP^) treated for 6 h with thapsigargin (Tg; 500 nM) or cycloheximide (CHX; 50 μg/mL).

The sensitivity of PRELID1 to translational attenuation suggests that this protein could similarly be reduced by PERK-regulated translational attenuation. As expected, PRELID1 was rapidly decreased in MEF^mtGFP^ cells treated with the ER stressor Tg (**Fig. 4C**). Tg-dependent reductions in PRELID1 were inhibited in cells deficient in *Yme1l*, indicating that YME1L was required for this process (**Fig. 4C**). Co-treatment with either the PERK kinase inhibitor GSK2656157 or the PERK signaling inhibitor ISRIB (**Fig. 1A**) blocked Tg-dependent reductions in PRELID1 (**Fig. 4D**). Similar results were observed for TIM17A. Further, Tg-dependent reductions in PRELID1 and TIM7A were also inhibited in *Perk^−/−^* MEFs (**Fig. 4E**). Reconstitution of *Perk^−/−^* with PERK^WT^, but not hypomorphic PERK^PSP^, restored Tg-induced degradation of these proteins. Importantly, CHX reduced PRELID1 and TIM17A in all genotypes, confirming that these proteins remain sensitive to translational attenuation even when PERK signaling is impaired (**Fig. 4E**). Interestingly, Tg-dependent reductions in PRELID1 were not inhibited in cells deficient in *Atf4* (**Fig. S4D**)^64^, a primary upstream transcription factor in the PERK pathway (**Fig. 1A**), indicating that this phenotype is independent of PERK-regulated transcriptional signaling. Collectively, these results suggest that PRELID1, like TIM17A^37^, is reduced during ER stress through a YME1L-dependent mechanism downstream of PERK-dependent translational attenuation.

### Depletion of PRELID1 rescues ER stress-dependent mitochondrial elongation in cells deficient in PERK signaling

Decreases in PRELID1 will slow trafficking of PA from the OMM to the IMM (**Fig. 4A**). This would increase PA on the OMM where it could promote mitochondrial elongation through the inhibition of DRP1. To define the importance of PERK-dependent reductions in PRELID1 on ER stress induced mitochondrial elongation, we used shRNA to deplete *Prelid1* from MEF^mtGFP^ cells and monitored mitochondrial morphology following treatment with Tg. We confirmed PRELID1 depletion by immunoblotting and demonstrated that loss of PRELID1 did not influence ER stress-induced, PERK-dependent mechanisms such as TIM17A degradation (**Fig. S5A**). *Prelid1* depletion did not basally influence mitochondrial morphology or inhibit Tg-induced mitochondrial elongation (**Fig. 5A,B**). This indicates that reduction of PRELID1, on its own, is not sufficient to increase mitochondrial elongation, likely reflecting the importance of PERK kinase-dependent increases in PA in this process.^52^ Consistent with this, co-treatment with the PERK kinase inhibitor GSK2656157 blocked Tg-induced mitochondrial elongation in *Prelid1*-deficient cells (**Fig. 5A,B**). However, PRELID1 depletion partially rescued mitochondrial elongation in cells co-treated with Tg and ISRIB – an inhibitor that blocks PERK signaling downstream of PERK kinase activity (**Fig. 1A**), but does not significantly inhibit ER stress dependent increases in mitochondrial PA (**Fig. S2D,E**). Identical results were observed in HeLa cells depleted of *PRELID1* and treated with Tg, GSK2656157, or ISRIB (**Fig. 5C,D**, **Fig. S5B**). These results implicate YME1L-dependent reductions of PRELID1 in the PERK-regulated mitochondrial elongation induced during ER stress.

**Figure 5.**
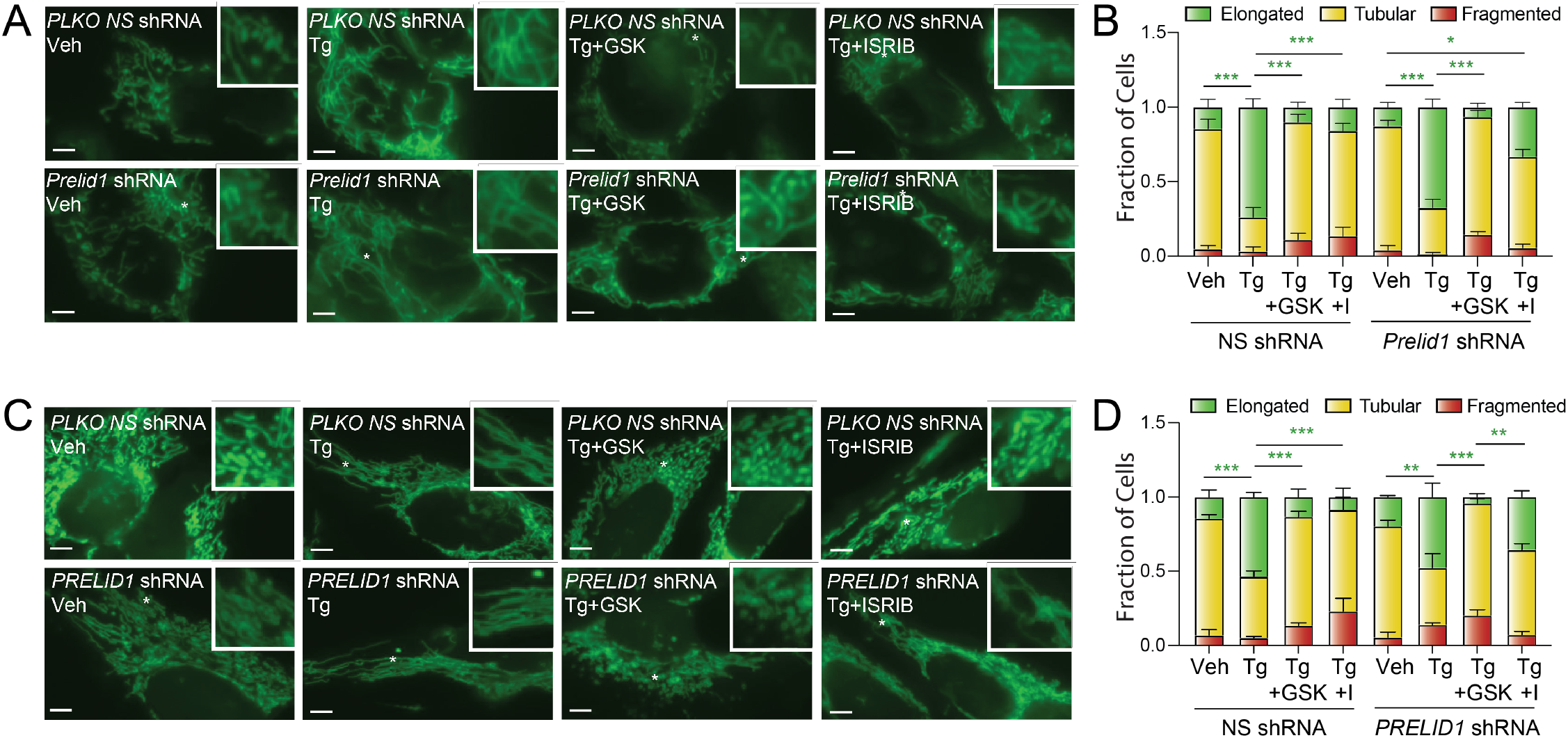
Reductions in PRELID1 contribute to ER stress induced mitochondrial elongation. **A.** Representative images of MEF^mtGFP^ cells expressing non-silencing (NS) or *Prelid1* shRNA treated for 3 h with thapsigargin (Tg; 500 nM), GSK265615 (GSK; 1 μM), and ISRIB (0.2 μM), as indicated. The inset shows 2-fold magnification of the image centered on the asterisk. Scale bars, 5 μm. **B.** Quantification of fragmented (red), tubular (yellow), or elongated (green) mitochondria from the images shown in (**A**). Error bars show SEM for n=5 experiments. *p<0.05; ***p<0.005 for 2-way ANOVA (green indicates comparisons between elongated mitochondria fractions). **C.** Representative images of HeLa cells expressing ^mt^GFP and non-silencing (NS) or *Prelid1* shRNA treated for 3 h with thapsigargin (Tg; 500 nM), GSK265615 (GSK; 1 μM), and ISRIB (0.2 μM), as indicated. The inset shows 2-fold magnification of the image centered on the asterisk. Scale bars, 5 μm. **D.** Quantification of fragmented (red), tubular (yellow), or elongated (green) mitochondria from the images shown in (**C**). Error bars show SEM for n=3 experiments. p-value reflects comparisons of elongated (green) mitochondria populations for the indicated conditions. **p<0.01; ***p<0.005 for 2-way ANOVA (green indicates comparisons between elongated mitochondria fractions).

## DISCUSSION

Mitochondrial elongation is an adaptive mechanism that protects mitochondria in response to diverse pathologic insults.^38,65–71^ Interestingly, numerous different mechanisms have been shown to promote mitochondrial elongation in response to different types of stress. For example, the accumulation of lysophosphatidic acid (LPA) on the outer mitochondrial membrane increases mitochondrial elongation through a MTCH2-dependent mechanism during starvation.^69^ Alternatively, HDAC6-dependent deacetylation of pro-fusion GTPase MFN1 increases mitochondrial length during glucose deprivation by enhancing the activity of organellar fusion pathways.^70^ Further, PKA-dependent phosphorylation or PARKIN-dependent ubiquitination of the pro-fission GTPase DRP1 inhibits mitochondrial fission and promotes mitochondrial elongation under a variety of different conditions.^71–73^ Despite these differences in mechanism, mitochondrial elongation similarly functions to prevent premature fragmentation, regulate mitochondria respiratory chain activity, and promote cell survival in response to diverse pathologic insults, including ER stress.^38,65–73^

ER stress promotes mitochondrial elongation through a process regulated by the PERK arm of the UPR.^38^ Here, we demonstrate that this PERK-dependent increase in elongated mitochondria is attributed to PERK-dependent remodeling of mitochondrial membrane PA (**Fig. 6**). Previously, ER stress was shown to increase cellular PA through a mechanism dependent on PERK kinase activity, but not eIF2α phosphorylation.^52^ This was suggested to involve direct, PERK-dependent phosphorylation of diacylglycerol (DAG; **Fig. 6**, step 1).^52^ Our results support the dependence of ER stress induced increases of PA on PERK kinase activity, showing that the PERK kinase inhibitor GSK2656157 reduces Tg-dependent increases of PA more efficiently than ISRIB, a compound that inhibits PERK signaling downstream of PERK kinase activity (**Fig. 1A**).

**Figure 6.**
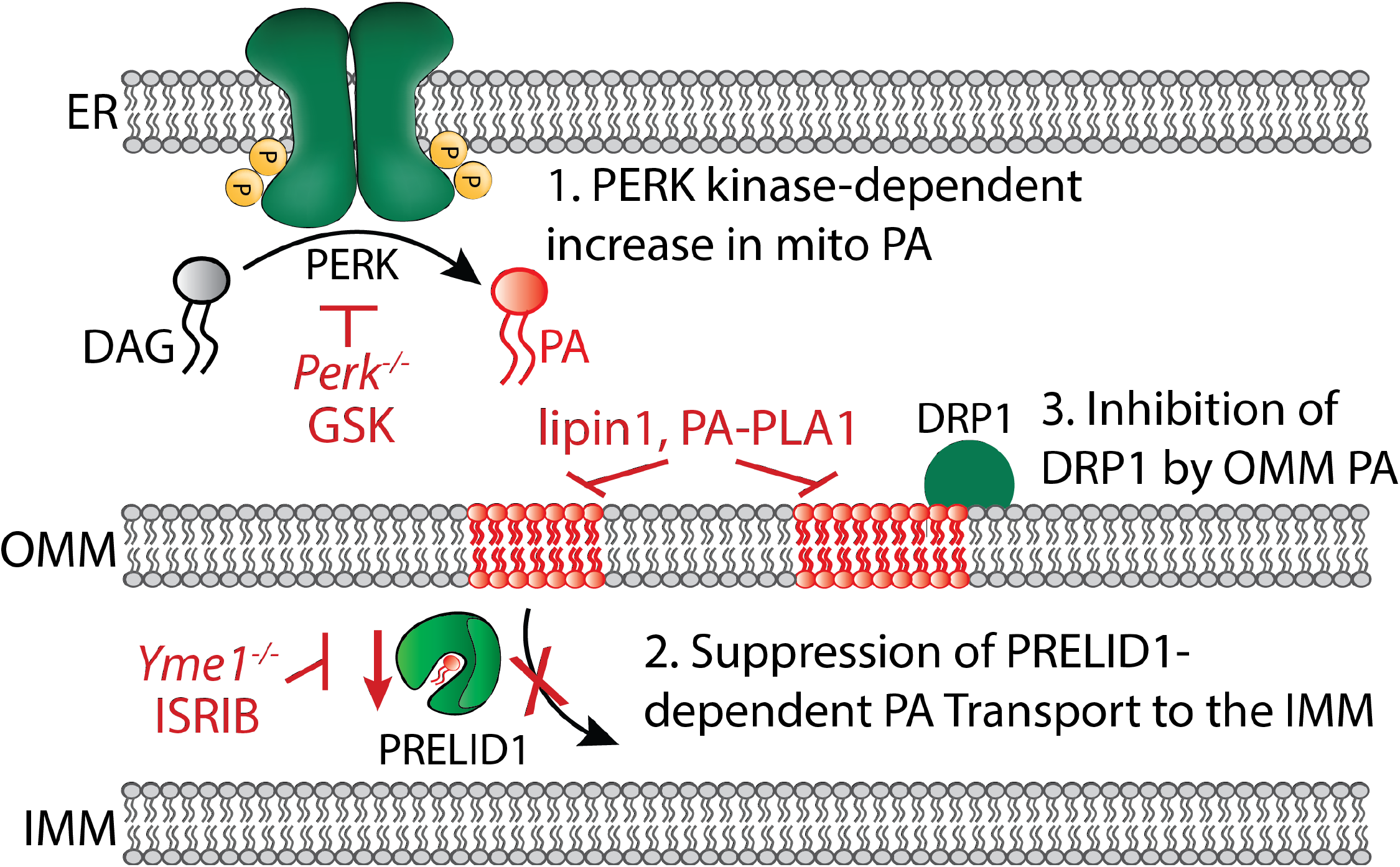
Proposed mechanism for PERK-regulated mitochondrial elongation during ER stress. In response to ER stress, PERK is activated, resulting in an increase in total and mitochondrial PA that was previously suggested to result from PERK kinase-dependent phosphorylation of diacylglycerol (DAG) to produce PA.^52^ YME1L degrades the intramitochondrial PA transporter PRELID1 downstream of PERK-dependent translational attenuation, limiting the trafficking of PA to the inner mitochondrial membrane (IMM) and resulting in the accumulation of PA on the outer mitochondrial membrane (OMM). This increase in OMM PA then promotes mitochondrial elongation by inhibiting mitochondrial fission, likely through the previously reported direct inhibition of DRP1 activity.^51^ Specific manipulations employed to manipulate specific steps in this pathway are shown.

However, our findings also suggest that PERK-dependent increases in PA are insufficient to drive the full extent of mitochondrial elongation observed during ER stress. To account for this, we show that ER stress-dependent increases in mitochondrial elongation also involve reductions in the intra-mitochondrial PA trafficking protein PRELID1 (**Fig. 6**, step 2). Consistent with previous results showing that PERK-dependent mitochondrial elongation requires PERK-dependent translational attenuation^38^, we show that PRELID1 is a short-lived mitochondrial protein that is degraded through a YME1L-dependent mechanism downstream of eIF2α phosphorylation-dependent reductions in translation. Intriguingly, we implicated PERK-dependent reductions in PRELID1 in ER stress induced mitochondrial elongation by showing that genetic depletion of PRELID1 partially rescues mitochondrial elongation in cells co-treated with the PERK signaling inhibitor ISRIB, but not the PERK kinase inhibitor GSK2656157. This highlights an important role for reductions in PRELID1, induced downstream of PERK-dependent translational attenuation through a YME1L-dependent mechanism, in the adaptive remodeling of mitochondrial membrane PA observed during ER stress.

The combination of PERK-dependent increases in total PA and YME1L-dependent decreases of PRELID1 should increase PA on the OMM during conditions of ER stress. Previous studies have shown that increases in OMM PA promote mitochondrial elongation through multiple mechanisms including direct inhibition of the pro-fission GTPase DRP1 (**Fig. 6**, step 3).^51^ Consistent with an important role for OMM PA in ER stress induced mitochondrial elongation, we show that overexpression of two different cytosolic PA lipases, Lipin1 and PA-PLA1, block mitochondrial elongation observed in Tg-treated cells. Further, we demonstrate that pre-treatment with the ER stressor Tg inhibits DRP1-dependent mitochondrial fragmentation in ionomycin-treated cells. Collectively, these results support a model whereby PERK-dependent increases in OMM PA promote mitochondrial elongation through a mechanism involving reductions in mitochondrial fission, likely mediated through the direct inhibition of the pro-fission GTPase DRP1 (**Fig. 6**).

PERK-regulated translational and transcriptional signaling regulate diverse aspects of mitochondrial proteostasis and function. Our results provide new insights into PERK-dependent remodeling of mitochondria by demonstrating that signaling through this UPR pathway promotes adaptive remodeling of mitochondrial membrane PA to induce protective organelle elongation during ER stress. As we and others continue studying the impact of PERK signaling on mitochondrial biology, additional mitochondrial pathways regulated through PERK signaling will also likely be identified, further expanding our understanding of the critical role for this stress-responsive signaling pathway in adapting mitochondria during ER stress. Moving forward, it will also be interesting to define how different PERK-dependent mitochondrial adaptations integrate to influence other aspects of mitochondrial function during conditions of stress. For example, recent cryo-electron tomography results indicate that mitochondrial elongation correlates with cristae remodeling in ER stressed cells, suggesting that these changes to bulk mitochondrial morphology and ultrastructure may be coordinated.^74^ Further, PRELID1-dependent trafficking of PA to the IMM is a critical step in the biosynthesis of cardiolipins – a mitochondrial-specific phospholipid important for regulating membrane morphology and multiple mitochondrial functions including respiratory chain activity and apoptotic signaling.^48^ While acute ER insults are unlikely to significantly influence cardiolipin levels owing to the higher stability of this phospholipid as compared to other membrane phospholipids^75^, chronic PERK-dependent translational attenuation, and subsequent sustained reductions in PRELID1, could lead to reductions in cardiolipin that impact mitochondrial energy metabolism and cell fate. Thus, an improved understanding of how different PERK-dependent alterations to mitochondrial morphology and function integrate will provide additional insight to the critical importance of this pathway in regulating mitochondria during conditions of ER stress.

The global importance of PERK in adapting mitochondria during ER stress also suggests that disruptions in this signaling could exacerbate mitochondrial dysfunction in disease. Genetic mutations in *EIF2AK3*, the gene that encodes PERK, are causatively associated with Wolcott-Rallison syndrome – a devastating disease characterized by early onset diabetes, skeletal deformities, and growth impairments.^76^ Further, a hypomorphic PERK haplotype is associated with tauopathies including progressive supranuclear palsy (PSP) and late stage Alzheimer’s disease.^15,40,41^ Interestingly, mitochondrial dysfunction has been implicated in all these disorders, suggesting that failure of PERK-dependent mitochondrial regulation is a contributing factor in disease pathogenesis. Consistent with this, we show that hypomorphic PSP-associated PERK alleles disrupt PERK-dependent mitochondrial elongation and YME1L-dependent PRELID1 degradation. In contrast, chronic PERK activation is also implicated in the pathogenesis of numerous neurodegenerative diseases involving mitochondrial dysfunction such as AD and prion disease.^77–81^ While the specific importance of PERK signaling on mitochondrial function in these diseases remains largely undefined, this suggests that PERK signaling, while adaptive during acute ER insults, could become detrimental to mitochondria in response to chronic ER insults potentially through mechanisms such as reductions in cardiolipin. Further investigations will be required to determine the specific impact of altered PERK signaling on mitochondria morphology and function in the context of these diseases to reveal both the pathologic and potentially therapeutic implications of PERK activity on the mitochondrial dysfunction observed in the pathogenesis of these disorders.

## MATERIALS AND METHODS

### Cell Culture, Transfections, Lentiviral Transduction, and CRISPR deletion

MEF^mtGFP^ (a kind gift from Peter Schultz, The Scripps Research Institute; TSRI)^82^, *Perk^+/+^ and Perk^−/−^* MEFs^24^, *Atf4^+/+^* and *Atf4^−/−^* MEFs^64^, HeLa cells (purchased from the ATCC), or HEK293T cells were cultured in DMEM (Corning-Cellgro) supplemented with 10% fetal bovine serum (FBS; Omega Scientific), 2 mM L-glutamine (GIBCO), 100 U/mL penicillin, and 100 mg/mL streptomycin (GIBCO). Cells were maintained at 37°C and 5% CO2. Non-essential amino acids (GIBCO) and 2-mercaptoethanol were also added to culture media of *Atf4^+/+^* and *Atf4^−/−^* MEFs and *Perk^+/+^* and *Perk^+/+^* MEFs. HeLa cells were transfected by calcium phosphate precipitation, as previously described.^38^ MEF cells were transfected with MEF Avalanche Transfection Reagent (EZ Biosystems) according to the manufacturer’s protocol. Lentivirus were prepared by transfecting HEK293T cells with pRSV-rev (Addgene #12253), pMDL-RRE (Addgene, #12251), pMD2.6 (Addgene #12259), and with the indicated shRNA in the pLKO.1 vector (Sigma) using calcium phosphate precipitation. After 24 h, the transfection media was removed and replaced with complete DMEM and incubated overnight for viral production. Virus containing media was removed the following day and filtered with a 0.45 μm syringe filter (Genessee Scientific). Polybrene (Sciquest/Fisher) was added to the viral containing media at a concentration of 10 μg/mL and the media was then added to HeLa or MEF^mtGFP^ cells. Stable pools of cells expressing non-silencing or gene-specific knockdowns were then generated through selection with puromycin (3 mg/mL for MEF cells and 1 mg/mL for HeLa). Knockdown was then confirmed by immunoblotting. *Yme1l* was deleted from MEF^mtGFP^ cells using CRISPR/Cas9. Briefly, cells were transfected with plasmid pSpCas9(BB)-2A-Puro (PX459; Addgene, #62988) containing sgRNA against *Yme1l* (GATCCAATATGAGATGTATGCCAAC AAACGTTGGCATACATCTCATATT) using MEF Avalanche, following manufacturers protocols. After transfection, cells were selected with puromycin and single clones were screened for YME1L depletion by qPCR and immunoblotting.

### Plasmids, shRNAs, and compounds

HA-LIPIN1^WT^, HA-LIPIN^Mut^, and mitoPLD-GFP overexpression constructs were kind gifts from Hiromi Sesaki (Johns Hopkins) and described previously.^51^ The PA-PLA1-GFP overexpression construct was from Addgene (#162880). *Perk^WT^* and *Perk^PSP^* overexpression plasmids were kind gifts from Jonathan Lin (Stanford). Plasmids containing shRNA were purchased from Sigma in the pLKO.1 vector: mouse *Prelid1* shRNA (TRCN0000345802), human *PRELID1* shRNA (TRCN0000130829). All compounds used in this study were purchased: thapsigargin (Tg; Fisher Scientific), GSK2656157 (BioVision Inc.), ISRIB (Sigma), CCCP (Sigma), and ionomycin (Sigma).

### Fluorescence Microscopy

HeLa cells transfected with ^mt^GFP or MEF^mtGFP^ cells were seeded at a density of 100,000 cells/well on glass-bottom dishes (MatTek) coated with poly-D-lysine (Sigma) or rat tail collagen 1 (GIBCO). Cells were then treated as indicated and images were recorded with an Olympus IX71 microscope with 60x oil objective (Olympus), a Hamamatsu C8484 camera (Hamamatsu Photonics), and HCI image software (Hamamatsu Photonics). At least 20 cells were imaged per condition for each experiment for quantification. Quantification was performed by blinding the images and then scoring cells based on the presence of primarily fragmented, tubular, or elongated mitochondria, as before^38^. Three different researchers scored each set of images and these scores were averaged for each individual experiment. All quantifications shown were performed for at least 3 independent experiments, where averages in morphology quantified from each individual experiment were then combined. The data were then prepared in PRISM (GraphPad, San Diego, CA) and plotted on a stacked bar plot to show the average morphology and standard error of the mean across all experiments. Statistical comparisons were performed using a 2-way ANOVA in PRISM, comparing the relative amounts of fragmented, tubular, or elongated mitochondria across different conditions.

### Phospholipid Quantification

For MS analysis of PA, whole cell pellets were resuspended in 500 μl of a cold hypotonic buffer consisting of 1 mM PBS, pH 7.4. The material was then homogenized on ice using a glass Dounce homogenizer (30 strokes). The homogenized sample was centrifuged at 500g for 4 min then the supernatant was transferred to a 1.5-ml microfuge tube and lyophilized overnight. The lyophilized material was weighed and normalized by total mass prior to performing a modified Bligh and Dyer extraction.^83^ The proceeding steps were done with glass pipettes and tubes to avoid plastic contamination. PA was extracted by the addition of 100 μl/mg of cold methanol containing an internal PA standard (Splash Lipidomix, Avanti) at a dilution of (1:50), followed by 50 μl/mg of cold chloroform (CHCl_3_) with occasional vortex mixing. Milli-Q H_2_0 containing 5 mM erythorbate was added at a volume of 50 μl /mg. The sample was agitated and centrifuged in glass test tubes at 200g for 10 min. The bottom phase was collected in a clean test tube, while the upper phase was re-extracted two additional times with methanol:CHCl_3_ (1:1, v:v) containing HCL at final concentration of 10 mM. The organic phases were combined and dried under vacuum to afford a lipid film that was stored at −80°C until MS analysis. Mitochondria enriched fractions were processed similar to whole cell samples except they were normalized via protein concentration as determined by the Pierce™ BCA Protein Assay kit (Thermo Scientific). In Brief, PA was extracted from the mitochondria enriched fractions using 10 μl/μg methanol, 5 μl/μg CHCl_3_ and 5 μl/μg milli-Q H_2_0 containing 5 mM erythorbate as previously described above.

Extracted lipid samples and a PA external standard (SPLASH Lipidomix, Avanti) were removed from the −80° C freezer and resuspended in 100 μl of methanol. Negative mode LC-MS analysis was performed on an Agilent 6230 ESI-TOF-MS System calibrated with a reference solution at m/z 1033.9881. A XBridge BEH C8 XP Column (Waters, 2.5 μm, 4.6 mm X 150 mm) was used at a flow rate of 500 μL/min, employing the following gradient: 30 to 100% solvent B over 30 min, 100% isocratic B for 10 min followed by a return to 30% B for 5 min. Solvents A consisted of MilliQ water and methanol (9:1, v:v). Solvents B consisted of acetonitrile: 2-propanol (5:3, v:v), and both contained 10 mM Piperidine, 10 mM ammonium acetate and 0.1% formic acid. Prior to processing, raw.d files were converted to the open format mzXML using MSConvert software, which is part of the ProteoWizard software toolkit.^84^ Mass detection was achieved using mzmine 2 wavelet algorithm, ADAP chromatogram builder and ADAP deconvolution^85^, which are part of the MZmine 2 software package.^86^ Initial lipid identifications were achieved using lipidmaps database with a m/z tolerance of 10 ppm, subsequently PA detected lipids were filtered out for further processing. These putative peaks were validated by aligning RT to the internal and external standards followed by graphical identification of PA lipids by plotting the Kendrick mass defect plot employing CH_2_ as the repeating unit. The quantification of all PA lipid class was normalized based on the abundance of the internal standard PA (15:0-18:1-d7-PA), which factors in extraction efficiency and sample handling. Total PA levels were then normalized to vehicle for the indicated number of independent experiments.

For quantification of PA by ELISA, MEF or HeLa cells were treated as indicated and collected on ice and then lysed with 20 mM Hepes pH 7.4, 100 mM NaCal, 1 mM EDTA, 1% Triton X100) supplemented with Pierce protease inhibitor (ThermoFisher). Protein concentrations for each sample were then quantified using the Bio-Rad Bradford assay. PA was then measured using the Human Phosphatidic Acid Antibody (IgM ELISA Kit (Lifeome Bioloabs) following the manufacturers protocol and monitoring fluorescence on a Tecan F250Pro microplate reader (Tecan).

For quantification of PA by fluorometric biochemical assay, MEF or HeLa cells were treated as indicated and collected on ice and then centrifuged and washed with cold PBS three times. Samples were then sonicated using the Misonix S-4000 sonicator then processed and PA was measured according the manufacturer’s protocol for the Total Phosphatidic Assay Kit (CellBio Labs).

### Immunoblotting and Antibodies

Whole cells were lysed at room temperature in HEPES lysis buffer (20 mM Hepes pH 7.4, 100 mM NaCal, 1 mM EDTA, 1% Triton X100) supplemented with 1x Pierce protease inhibitor (ThermoFisher). Total protein concentrations of lysates were then normalized using the Bio-Rad protein assay and lystaes were combined with 1x Laemmli buffer supplemented with 100 mM DTT and boiled for 5 min. Samples were then separated by SDS-PAGE and transferred to nitrocellulose membranes (Bio-Rad). Membranes were then blocked with 5% milk in tris-buffered saline (TB) and then incubated overnight at 4°C with the indicated primary antibody. The next day, membranes were washed in TBS supplemented with Tween, incubated with the species appropriate secondary antibody conjugated to IR-Dye (LICOR Biosciences) and then imaged using an Odyssey Infrared Imaging System (LICOR Biosciences). Quantification was then carried out using the LICOR Imaging Studio software. Primary antibodies were acquired from commercial sources and used in the indicated dilutions in antibody buffer (50mM Tris [pH 7.5], 150mM NaCl supplemented with 5% BSA (w/v) and 0.1% *NaN*_3_(w/v)): Tim17a (Thermo Scientific, 1:1000), PRELID1 [aa27-54] (Sciquest, 1:1000), YME1L (Abgent, 1:1000), ATF4 (Cell Signaling, 1:500), Tubulin [B-5-1-2] (Sigma, 1:5000), Tim23 (BD Transduction Labs, 1:1000), HSP60 [LK1] (Thermo Scientific, 1:1000), PERK (Sciquest, 1:1000), HA [Clone: 16B12] (Biolegend, 1: 1000), GFP (B2) (Sciquest, 1:1000), OPA1 (BD Transduction Labs, 1:2000).

### Quantitative Polymerase Chain Reaction (qPCR)

The relative mRNA expression of target genes was measured using quantitative RT-PCR. Cells were treated as indicated and then washed with phosphate buffered saline (PBD; Gibco). RNA was extracted using Quick-RNA MiniPrepKit (Zymo Research) according to the manufacturers protocol. RNA (500 ng) was then converted to cDNA using the QuantiTect Reverse Transcription Kit (Qiagen). qPCR reactions were prepared using Power SYBR Green PCR Master Mix (Applied Biosystems), and primers (below) were obtained from Integrated DNA Technologies. Amplification reactions were run in an ABI 7900HT Fast Real Time PCR machine with an initial melting period of 95 °C for 5 min and then 45 cycles of 10 s at 95 °C, 30 s at 60 °C.

#### Primers used in this study

*Human ASNS:* forward: GCAGCTGAAAGAAGCCCAAG; reverse: AGCCTGAATGCCTTCCTCAG

*Human CHOP/DDI3:* forward: ACCAAGGGAGAACCAGGAAACG; reverse TCACCATTCGGTCAATCAGAGC

*Human HSPA5/BIP*: forward: GCCTGTATTTCTAGACCTGCC; reverse, TTCATCTTGCCAGCCAGTTG

*Human RIBOP:* forward: CGT CGC CTC CTA CCT GCT; reverse, CCA TTC AGC TCA CTG ATA ACC TTG

### Membrane Depolarization

MEFs were seeded at a density of 85,000 cells/well of a 6-well plate and treated with 500 nM Tg for 3h prior to collection. CCCP (10 μM) was added 50 min before collection, followed by 200 nM TMRE (Thermofisher) 20 min before collection. Samples were collected using TrypLE Express and cell culture media. Following a brief centrifugation, cell pellets were washed in DPBS (Gibco) and resuspended in DPBS supplemented with 5% BSA. Fluorescence intensity of TMRE for 20,000 cells/condition was recorded on the PE channel of a BD Biosciences LSR II analytical flow cytometer. Data are presented as geometric mean of the fluorescence intensity from three experiments normalized to vehicle-treated cells.

### Statistical Analysis

Statistics were calculated in PRISM 9 (GraphPad, San Diego, CA). Data are presented as mean +/− SEM and were analyzed by 2-way ANOVA with Tukey’s multiple correction test or the appropriate student t-tests, as indicated in the accompanying figure legends. Indications of non-significant interactions from 2-way ANOVA were generally omitted for clarity.

## ACKNOWLEDGEMENTS.

We thank Jessica Rosarda and Cristina Puchades for experiments and analysis related to this work. We thank Benjamin Barad, Michaela Medina, Enrique Saez, and Katja Lamia for critical reading of this manuscript. This work was funded by the National Institutes of Health (NS095892 to RLW), a National Science Foundation predoctoral fellowship (VP), and the George E. Hewitt Postdoctoral fellowship (VD).

## AUTHOR CONTRIBUTIONS.

VP, CC, and JL conceived, designed, performed, and interpreted the experiments. VD, KB, and AM performed experiments. DG and RLW interpreted experiments. RLW supervised the project. VP and RLW wrote the manuscript. VP, CC, JL, VD, KB, JWK, DG, and RLW provided critical revisions to the manuscript and approval for submission.

---

**Figure S1.**
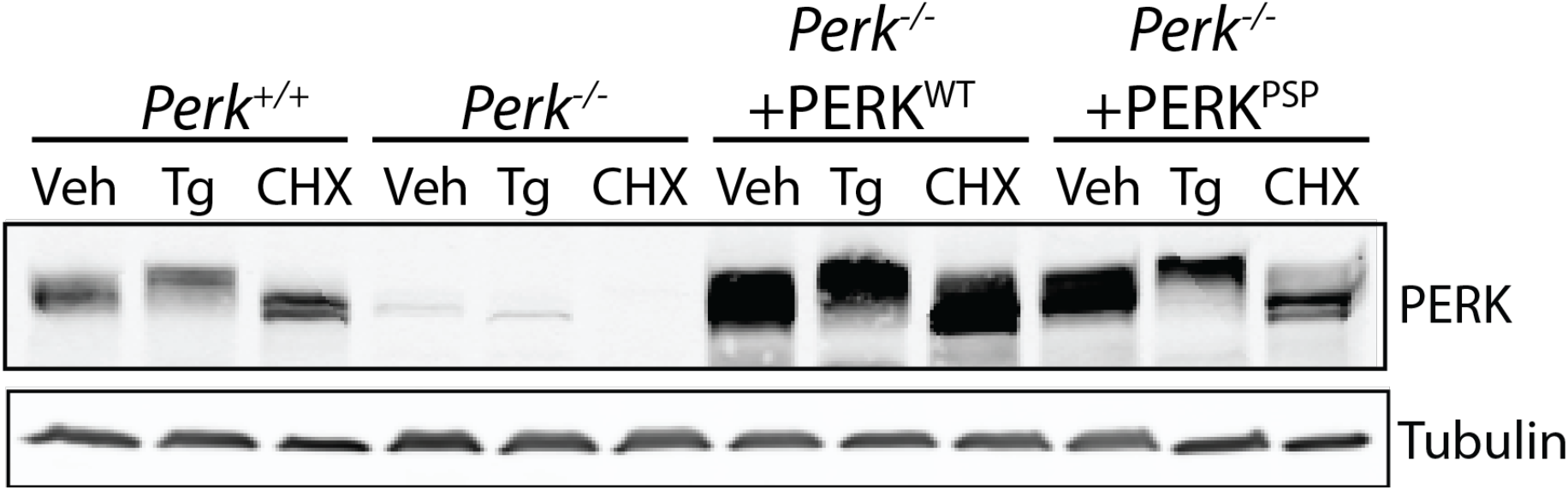
(Supplement to Figure 1. ER stress induced mitochondrial elongation is impaired in cells expressing hypomorphic PERK variants). Immunoblot of lysates prepared from *Perk^+/+^* MEFs, *Perk^−/−^* MEFs, or *Perk^−/−^* MEFs transfected with wild-type PERK^WT^ or the PSP-associated PERK allele (PERK^PSP^) treated for 6 h with thapsigargin (Tg; 500 nM) or cycloheximide (CHX; 50 μg/mL).

**Figure S2.**
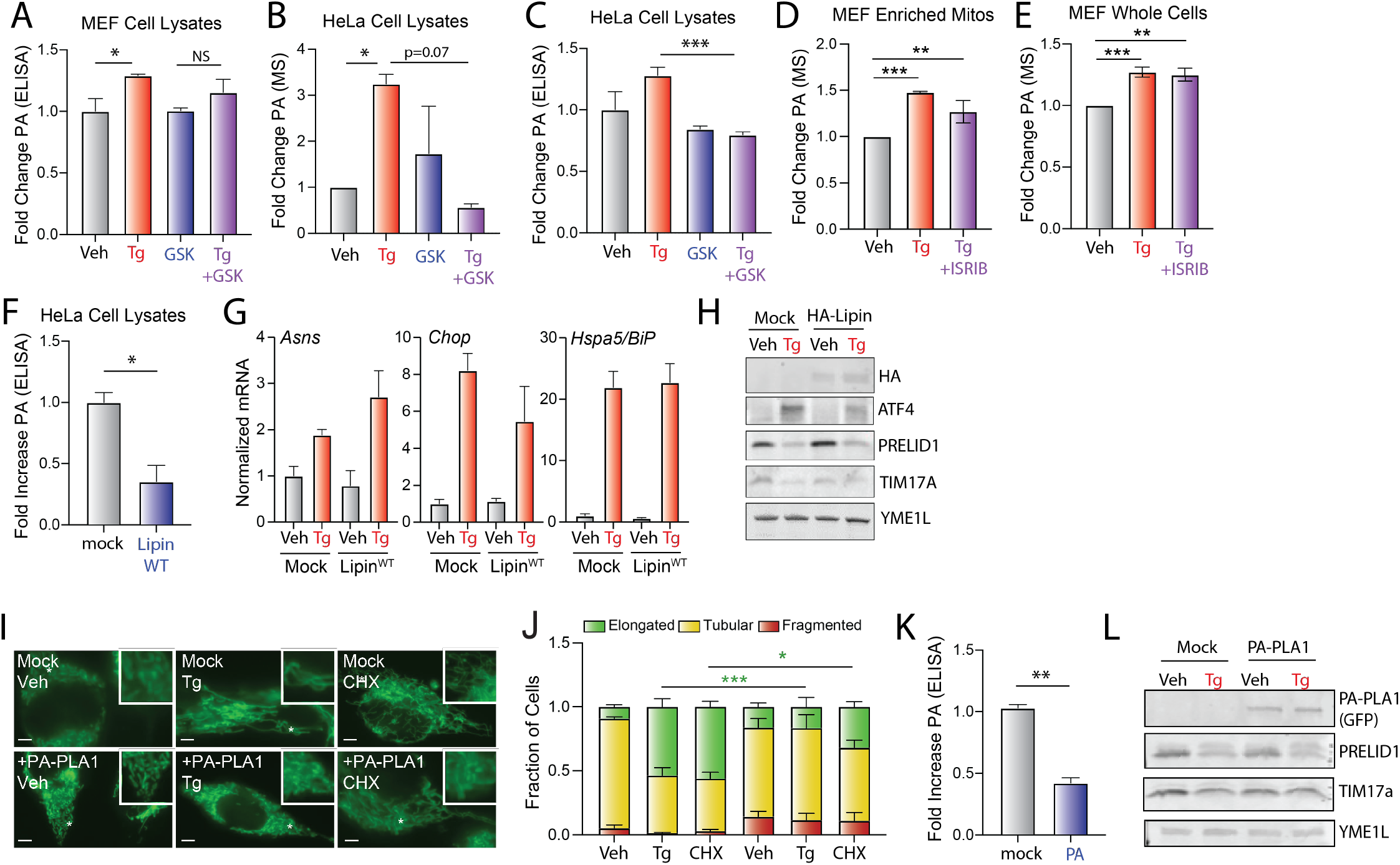
(Supplement to Figure 2. Overexpression of PA lipases inhibits ER stress induced mitochondrial elongation.). **A.** Relative phosphatidic acid (PA), measured by ELISA (see Materials and Methods), in whole cell extracts isolated from MEF cells treated for 3 h with vehicle, thapsigargin (Tg, 500 nM), and/or GSK265615 (1 μM). Error bars show SEM for n=3 replicates *p<0.05 for unpaired t-test. **B,C**. Relative PA, measured by mass spectrometry (**B**) or ELISA (**C**), in lysates prepared from HeLa cells treated for 3 h with vehicle, thapsigargin (Tg, 500 nM), and/or GSK265615 (1 μM). Error bars show SEM for n=3 replicates *p<0.05 for paired t-test in (**B**) and ***p<0.005 for unpaired t-test in (**C**). **D,E**. Relative PA levels, measured by mass spectrometry, in mitochondrial enriched fractions or whole lysates from MEF cells treated for 3 h with vehicle, thapsigargin (Tg; 500 nM), and ISRIB (0.2 μM). Error bars show SEM for n=3 replicates. **p<0.01, ***p<0.005 for paired t-test. **F**. Relative PA, measured by ELISA, in lysates of HeLa cells expressing mock or Lipin^WT^. Error bars show SEM for n=3 replicates. *p<0.05 for unpaired. **G**. Expression, measured by qPCR, of *Asns*, *Chop*, and *Hspa5/BiP* in HeLa cells expressing mock or Lipin^WT^ treated for 3 h with vehicle or thapsigargin (Tg, 500 nM). **H.** Immunoblot of lysates prepared form HeLa cells expressing mock or Lipin^WT^ treated for 3 h with vehicle or thapsigargin (Tg; 500 nM). Note that the Lipin construct is HA tagged allowing detection with the HA antibody. **I.** Representative images of HeLa cells expressing ^mt^GFP transfected with mock or GFP-tagged PA-PLA1 then treated for 3 h with thapsigargin (Tg; 500 nM) or cycloheximide (CHX; 50 μg/mL). The inset shows 2-fold magnification of the image centered on the asterisk. Scale bars, 5 μm. Note that the presence of GFP on PA-PLA1 did not influence our ability to monitor mitochondrial morphology in these cells. **J.** Quantification of fragmented (red), tubular (yellow), or elongated (green) mitochondria from the images shown in (**I**). Error bars show SEM for n=3 experiments. *p<0.05, ***p<0.005 for 2-way ANOVA (green indicates comparisons between elongated mitochondria fractions). **K.** Relative PA, measured by ELISA, in lysates of HeLa cells expressing mock or PA-PLA1. Error bars show SEM for n=3 replicates. Error bars show SEM for n=3 replicates. **p<0.01 for unpaired t-test. **L.** Immunoblot of lysates prepared form HeLa cells expressing mock or PA-PLA1 treated for 3 h with vehicle or thapsigargin (Tg; 500 nM).

**Figure S3.**
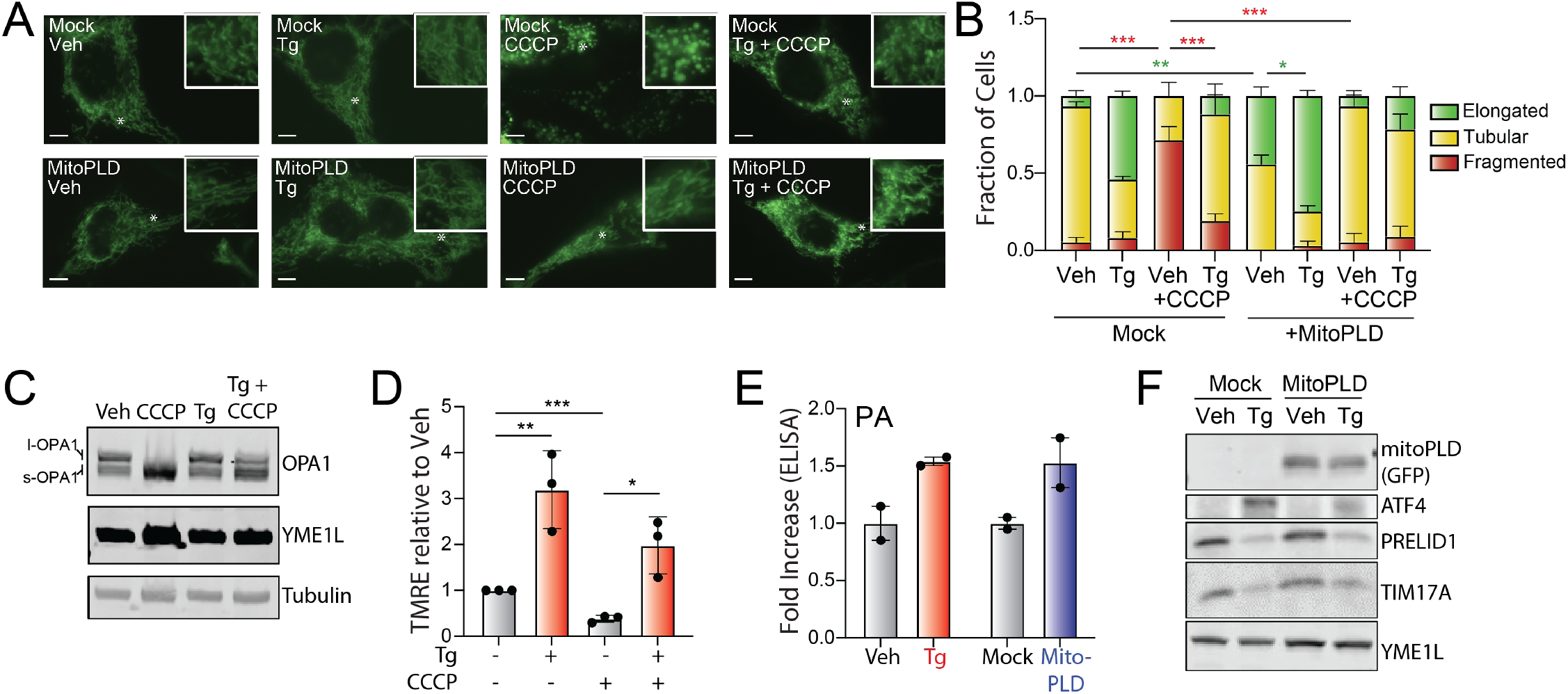
(Supplement to Figure 3. ER stress induced mitochondrial elongation inhibits Ionomycin-induced mitochondrial fragmentation.) **A.** Representative images of HeLa cells expressing ^mt^GFP transfected with mock or mitoPLD^GFP^ and then pretreated for 3 h with thapsigargin (Tg; 500 nM) followed by a 30 min incubation with CCCP (1 μM), as indicated. The inset shows 2-fold magnification of the image centered on the asterisk. Scale bars, 5 μm. Note that the expression of the mitoPLD^GFP^ did not impair our ability to monitor mitochondrial morphology in these cells. **B.** Quantification of fragmented (red), tubular (yellow), or elongated (green) mitochondria from the images shown in (**A**). Error bars show SEM for n=3 experiments. p-value reflects comparisons of elongated (green) or fragmented (red) mitochondria populations for the indicated conditions. *p<0.05, **p<0.01, ***p<0.005 for 2-way ANOVA (red indicates comparison between fragmented mitochondria fractions; green indicates comparisons between elongated mitochondria fractions). **C.** Immunoblot of lysates prepared from MEF^mtGFP^ cells pre-treated for 3 h with thapsigargin (Tg; 500 nM) then challenged with CCCP for 30 min. **D.** Mitochondrial polarization, measured by TMRE fluorescence, in MEF cells pre-treated for 3 h with thapisgargin (Tg; 500 nM) then challenged for 30 min with CCCP (10 μM). Error bars show SEM for n=3 replicates. *p<0.05, **p<0.01, ***p<0.005 for paired t-tests. **E**. Phosphatidic acid (PA), measured by ELISA, in HeLa cells treated with thapsigargin (Tg; 500 nM, 3 h) or expressing mock or mitoPLD. Error bars show SEM for n=2 replicates. Individual replicates are shown. **F.** Immunoblot of lysates prepared om HeLa cells transfected with mock or mitoPLD and treated for 3 h with vehicle or thapsigargin (Tg; 500 nM). Note mitoPLD is tagged with GFP allowing detection of this protein with the GFP antibody.

**Figure S4.**
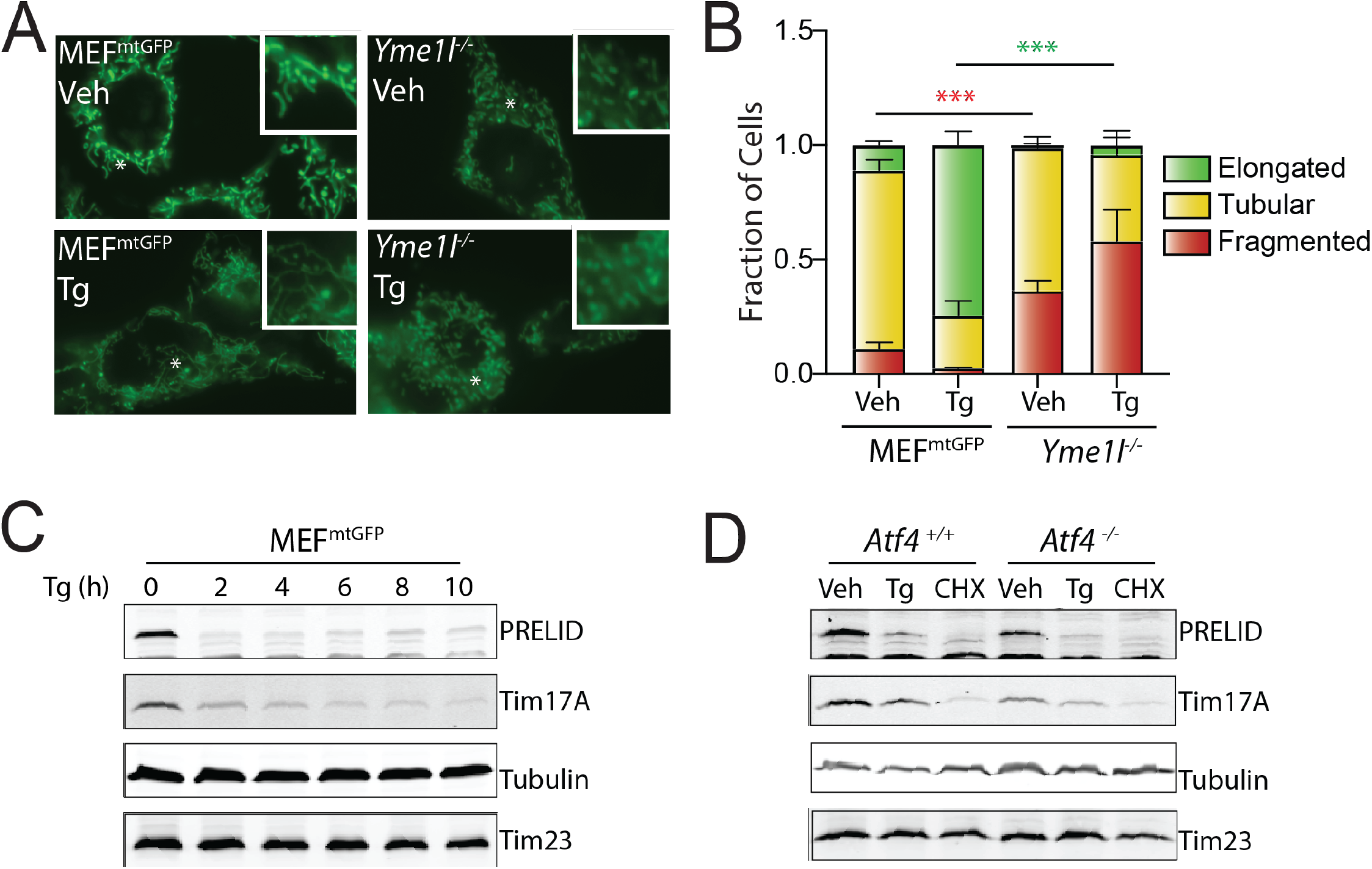
(Supplement to Figure 4. ER stress reduces PRELID1 through a YME1L-dependent mechanism downstream of PERK-dependent translational attenuation). **A.** Representative images of MEF^mtGFP^ cells and *Yme1l*-deficient MEF^mtGFP^ cells treated for 6 h with thapsigargin (Tg; 50 nM). The inset shows 2-fold magnification of the image centered on the asterisk. Scale bars, 5 μm. **B.** Quantification of fragmented (red), tubular (yellow), or elongated (green) mitochondria from the images shown in (**A**). Error bars show SEM for n=7 experiments. p-value reflects comparisons of elongated (green) or fragmented (red) mitochondria populations for the indicated conditions. ***p<0.005 for 2-way ANOVA (red indicates comparison between fragmented mitochondria fractions; green indicates comparisons between elongated mitochondria fractions). **C.** Immunoblot of lysates prepared from MEF^mtGFP^ cells treated with thapsigargin (Tg; 500 nM) for the indicated time. **D.** Immunoblot of lysates prepared from *Atf4^+/+^* and *Atf4^−/−^* MEFs treated with thapsigargin (Tg; 500 nM) or CHX (50 μg/mL) for 3 h.

**Figure S5.**
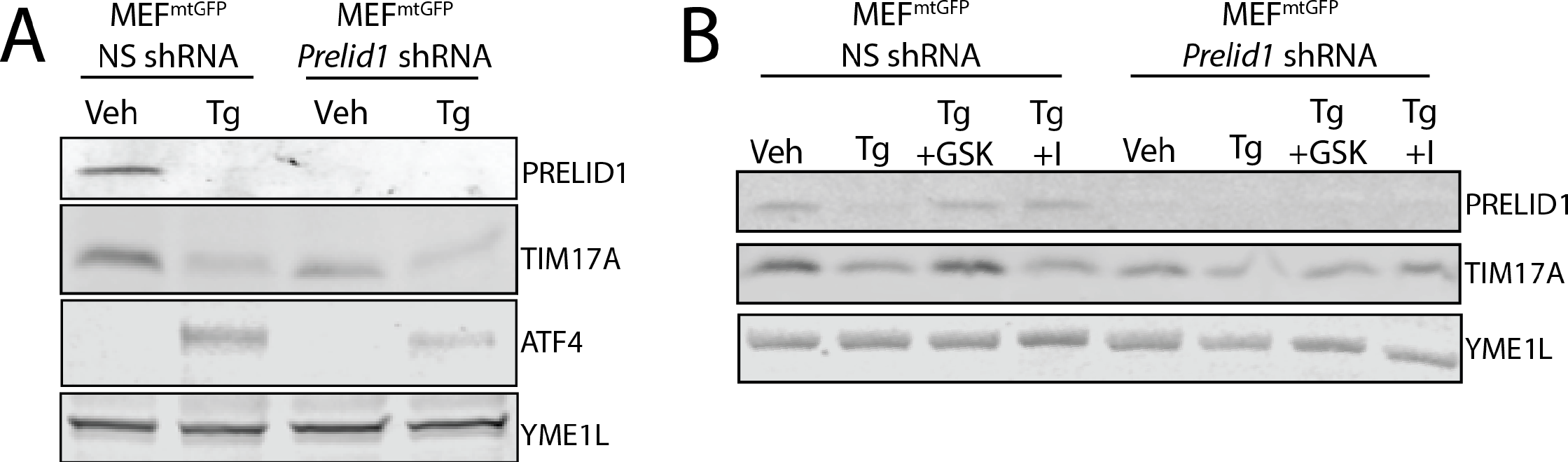
(Supplement to Figure 5. Reductions in PRELID1 contribute to ER stress induced mitochondrial elongation.) **A.** Immunoblot of lysates prepared from MEF^mtGFP^ expressing non-silencing (NS) or *Prelid1* shRNA and treated for 3 h with vehicle or thapsigargin (Tg, 500 nM). **B.** Immunoblot of lysates prepared from HeLa cells expressing non-silencing (NS) or *PRELID1* shRNA and treated for 3 h with vehicle or thapsigargin (Tg, 500 nM).

